# Olfactory fear conditioning non-specifically enhances glomerular odor responses and increases representational overlap of learned and neutral odors

**DOI:** 10.1101/194415

**Authors:** Jordan M. Ross, Max L. Fletcher

## Abstract

Associative fear learning produces fear toward the conditioned stimulus (CS) and often generalization, the expansion of fear from the CS to similar, unlearned stimuli. However, how fear learning affects early sensory processing of learned and unlearned stimuli in relation to behavioral fear responses to these stimuli remains unclear. We subjected mice to a classical olfactory fear conditioning paradigm and used awake, *in vivo* calcium imaging to quantify learning-induced changes in glomerular odor responses, which constitutes the first site of olfactory processing in the brain. The results demonstrate that olfactory fear learning non-specifically enhances glomerular odor representations in a learning-dependent manner and increases representational similarity between the CS and non-conditioned odors. This mechanism may prime the system towards generalization of learned fear. Additionally, CS-specific enhancements remain even when associative learning is blocked; suggesting two separate mechanisms lead to enhanced glomerular responses following odor-shock pairings.

## Introduction

Associative fear learning, in which an organism learns that a given stimulus predicts an aversive outcome, produces behavioral displays of fear upon subsequent encounters with that stimulus (CS). In addition, this form of learning often produces robust generalization, the expansion of fear from the threat-predictive CS to other, unlearned stimuli. Studies in different sensory systems demonstrate that fear learning alters the neural networks responsible for encoding the CS-fear association (Bakin and Weinberger, 1990; Grundemann and Luthi, 2015; Herry et al., 2008; Johansen et al., 2011; Maren, 2003a, b, 2005; Rogan et al., 1997; Sadrian and Wilson, 2015; Walker et al., 2005; Weinberger, 2007), but it remains unclear how representations of non-conditioned stimuli are altered, especially in relation to behavioral generalization. In the case of olfactory fear conditioning, in which an odor is paired with an aversive outcome, studies indicate altered CS encoding (Coopersmith et al., 1986; Fletcher, 2012; Freeman and Schneider, 1982; Kass et al., 2013). However, how early olfactory sensory processing changes as a result of simple fear learning and how this relates to generalization remains unknown.

Olfactory bulb (OB) glomeruli are the sites of synaptic contact between olfactory sensory neuron (OSN) axons and the dendrites of OB output cells, such as mitral/tufted (M/T) cells (Buck and Axel, 1991). Odor molecules bind to OSNs expressing the same receptor gene, which, in turn, project to only one or two OB glomeruli. This preferential receptor binding generates unique patterns of glomerular activation, which codes the experienced as a spatial map and by strength of glomerular responses (Bozza et al., 2004; Fletcher et al., 2009; Mori et al., 2006). This odor-specific coding constitutes the first site of OB processing in the central nervous system before it is projected to higher-order processing centers in the brain, including those involved in associative learning (Shipley and Ennis, 1996). One such region, the amygdala, is an integral component of the fear learning circuit that is involved in both acquisition and expression of fear memories (Herry and Johansen, 2014). While many posit learning-induced amygdalar hyperactivity as a putative cause of fear generalization (Morey et al., 2013; Shin et al., 2005), there is a paucity of data regarding the role sensory processing transformations may play in these behavioral outcomes.

Previous studies establish olfactory learning alters information processing at the glomerular layer (Buonviso and Chaput, 2000; Fletcher and Wilson, 2003; Nicol et al., 2014; Sullivan and Leon, 1986; Wilson and Leon, 1988; Wilson et al., 1985; Woo et al., 1987; Woo and Leon, 1991) however, recent technological advances have allowed investigation of learning-induced transformations of spatial representations of odors (Fletcher, 2012; Kass et al., 2013). While these studies clearly indicate learning alters glomerular representations of odorants, they investigated effects in anesthetized mice and recent studies demonstrate variable glomerular coding for the same odors between awake and anesthetized states (Blauvelt et al., 2013; Kato et al., 2012). Furthermore, studies have yet to investigate how learning to fear the conditioned odor might affect sensory processing of neutral, unlearned odorants, an important question in light of behavioral fear generalization. The extent to which olfactory aversive learning modulates sensory processing in awake mice and how such alterations might contribute to behavioral generalization remains unknown.

Using awake, behaving calcium imaging we report, for the first time, that a single day fear learning paradigm leads to long-lasting increased glomerular responses to not just the learned odor (CS) but also to other, non-conditioned odors. Such enhancements lead to increasingly overlapping glomerular representations between the conditioned and neutral odors. Furthermore, we demonstrate this global glomerular enhancement is dependent on amygdalar activation during acquisition but not expression, suggesting it requires associative fear learning and the alterations occur during or shortly after learning acquisition. Together, these results indicate that olfactory learning induces changes as early as the first synapse in the olfactory system and alters glomerular odor coding in a way that may provide a switch between generalized or specific behavioral outcomes of learning.

## Results

### Olfactory aversive conditioning produces fear generalization to multiple odors

To investigate the extent to which olfactory fear conditioning alters both behavioral and olfactory bulb (OB) glomerular responses in awake mice, we adapted a method for awake, behaving imaging using a treadmill that provides mice full control of their forward/reverse motion while still allowing for head-fixation under a microscope. We imaged glomerular activity in Thy1-GCaMP3 mice, where the post-synaptic OB excitatory cells express the fluorescent calcium indicator GCaMP3, to a panel of 5 odors. The panel comprised diverse odors to assess the extent of behavioral generalization as well as OB glomerular transformations. In addition to imaging OB activity in these mice, we also subjected each mouse to a behavioral paradigm.

We began by imaging glomerular responses to the panel of odors for two consecutive days, establishing baseline responses. After the second imaging session, each mouse was placed into one of three training conditions: Odor only (n = 5), Shock only (n = 5), or Paired (n = 5). Mice in the Paired condition received 6 odor-shock pairs (10s E5; 0.5s, 0.8mA; 3m ISI), mice in the Shock only condition received 6 unpaired foot shocks, and mice in the Odor only condition received 6 unpaired odor presentations. Approximately 24 hours after conditioning, mice were placed in a novel chamber and were tested for behavioral freezing in response to a presentation of each odor in the imaging panel (Figure 1A). Only mice in the Paired condition displayed conditioned freezing, with a main effect of stimulus on freezing (ANOVA: F_5, 24_ = 12.984, p < 0.0001, η^2^ = 0.730). Post hoc analysis indicates only baseline freezing was significantly different from freezing to the CS (E5), p < 0.0001, indicating the paired training paradigm produces strong fear learning and broad behavioral fear generalization across all odors (Figure 1D). There was no difference in behavioral freezing for Odor only mice (AVOVA: F_5, 24_ = 0.635, p = 0.675, η^2^ = 0.117) (Figure 1B); or Shock only mice across the different testing stimuli (ANOVA: F_5, 24_ = 1.933, p = 0.126, η^2^ = 0.287) (Figure 1C).

**Figure 1.**
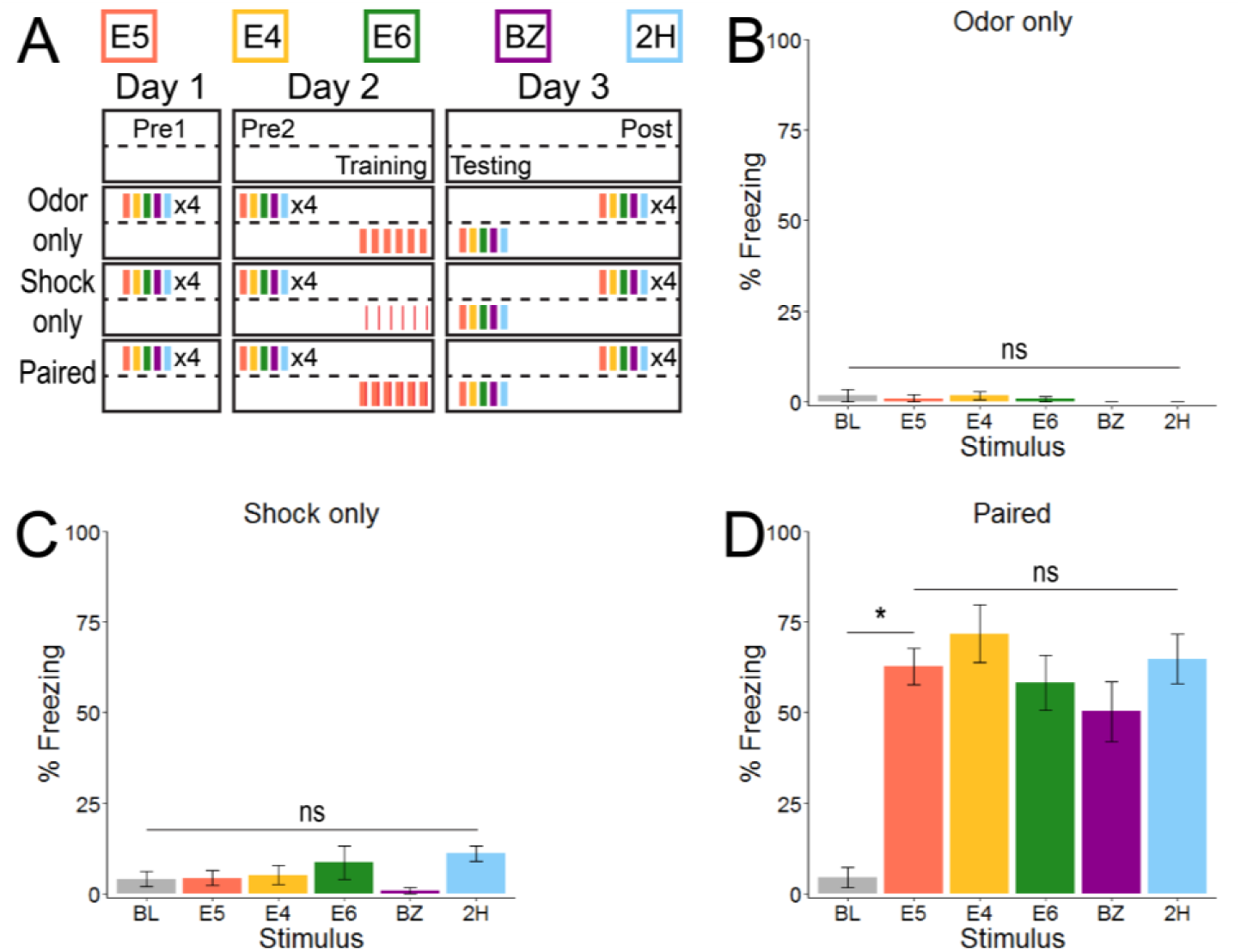
Olfactory aversive conditioning results in robust olfactory fear and generalization. (A) Timeline detailing the odors used (top) and both imaging (above dotted line) and behavioral (below dotted line) experiments for each group. (B-D) 24 hours after training, mice were exposed to each of the five odors (E5, E4, E6, BZ, & 2H) and freezing was measured. Odor only mice (B) and Shock only mice (C) did not learn to fear E5, as freezing was not significantly different than baseline (BL), and did not generalize freezing from E5 to other odors. Paired mice (D) froze significantly more to E5 than baseline, indicating acquired fear to the CS. Mice generalized fear across all other tested odors (freezing to other odors not significantly different from freezing to E5). Data presented mean ± SEM.

### Olfactory aversive conditioning non-specifically potentiates glomerular responses to all odors in awake mice

Following testing, mice underwent the final imaging session to assess the effect of conditioning on glomerular responses (Figure 2A-C). We first tested whether there was an effect of time (day) on overall glomerular response amplitude. There was a main effect of time on glomerular responses of Odor only mice (n = 570 averaged glomerular responses; RM ANOVA: F_1.930, 1098.288_ = 2636.434, p < 0.0001, η^2^ = 0.822), Shock only mice (n = 508 averaged glomerular responses; RM ANOVA: F_1.895, 960.989_ = 2665.88, p < 0.0001, η^2^ = 0.840), and Paired mice (n = 586 averaged glomerular responses; RM ANOVA: F_1.743, 1019.743_ = 907.598, p < 0.0001, η^2^ = 0.608). Post hoc analyses indicate that glomerular responses were significantly different at all time points for each of the three groups. The glomerular responses of Odor only and Shock only mice decreased from Pre1 to Pre2 (Figures 2A’, 2B’ and 3A, 3B) and were further reduced after training (Figures 2A”, 2B” and 3A, 3B), supporting the idea that glomerular responses decrease with increasing odor familiarity. Responses of Paired mice also decreased from Pre1 to Pre2 (Figures 2C’ and 3C), but were significantly enhanced after training (Figure 2C’’ and 3C).

**Figure 2.**
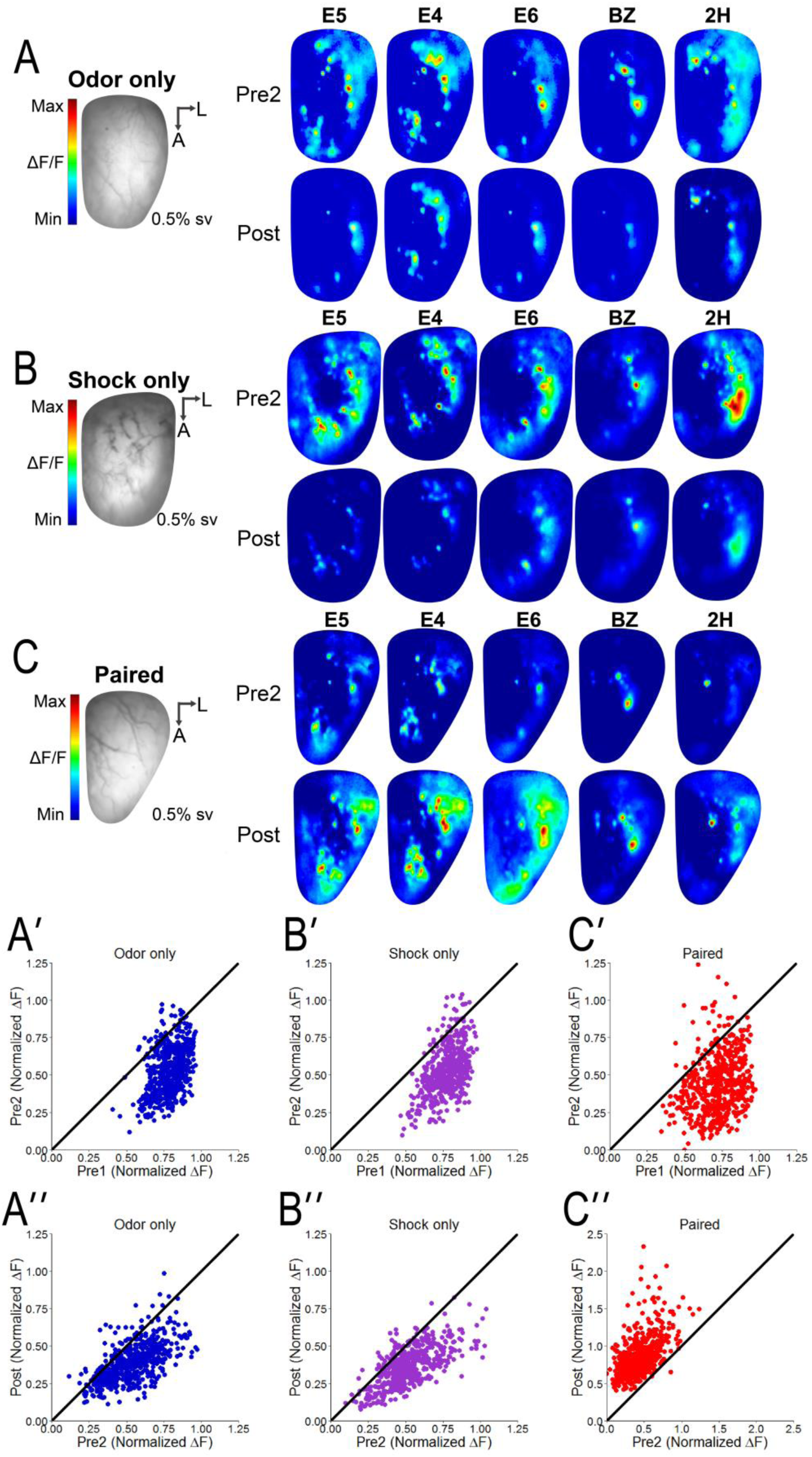
Olfactory aversive conditioning enhances glomerular responses. (A-C) Resting light intensity (RLI) frames and psuedocolored averaged Pre2 and Post glomerular response maps from representative Odor only (A), Shock only (B), and Paired (C) mice. (A’-C’) Scatterplots showing normalized Pre1 (x-axis) and corresponding normalized Pre2 (y-axis) values for each recorded glomerulus from all Odor only (A’), Shock only (B’), and Paired (C’) mice. (A’’-C’’) Scatterplots showing normalized Pre2 (x-axis) and corresponding normalized Post (y-axis) values for each recorded glomerulus from all Odor only (A’’), Shock only (B’’), and Paired (C’’) mice. Solid black line represents theoretical “no change” line. Glomerular responses generally decrease from Pre1 to Pre2 for all groups, as evidenced by the majority of points falling below the no change line (A’-C’). Glomerular responses also generally decrease from Pre2 to Post for Odor only (A’’) and Shock only (B’’) mice while almost all glomerular responses increase from Pre2 to Post for Paired mice (C’’), as evidenced by points falling above no change line.

As respiratory rate could affect signal magnitude, we examined whether the breathing (mean instantaneous frequency) might be responsible for the observed changes reported above. In order to assess mean instantaneous respiration frequency (MIF), we first extracted the respiratory signal from a fluorescent trace in at least 3 odor trials for each odor before (Pre2) and after (Post) training for all Odor only, Shock only, and Paired mice. Odor-evoked MIF was not significantly different between groups before or after training (ANOVA Pre2: F_2, 73_ = 2.300, p = 0.108; ANOVA Post: F_2, 73_ = 0.202, p = 0.817). Additionally, there was no effect of odor on odor-evoked MIF before or after training for any group (data not shown). During the post-training imaging session (Post), the difference between MIF before and after odor onset (as measured by calculating the MIF for all respirations before odor onset and the MIF of the first four respirations after odor onset) is not significantly different between groups (ANOVA F_2, 72_ = 0.207, p = 0.814). This indicates any recorded changes in the fluorescent signal is not due to differences in respiration between groups but reflects real changes in glomerular activity as a result of experimental condition.

We next examined whether the glomerular response changes were caused by a single odor by separately analyzing glomerular responses across time for each individual odor. Responses to all odors in the Odor only group exhibited similar decreases over time (RM ANOVA E5: n = 133, F_1.711, 225.815_ = 579.243, p < 0.0001, η^2^ = 0.814; E4: n = 136, F_2, 270_ = 1020.020, p < 0.0001, η2 = 0.883; E6: n = 82, F_2, 162_ = 507.476, p < 0.0001, η^2^ n = 118, F_1.597, 186.899_ = 1165.399, p < 0.0001, η^2^ = 0.909; 2H: n = 101, F_2, 200_ = 253.07, p < 0.0001, η2 = 0.717) and post hoc analyses indicate responses are significantly decreased from the preceding time point for all odors (Figure 3A). Analogous decreases across odors were observed in the responses from Shock only mice (RM ANOVA E5: n = 120, F_1.637, 194.820_ = 531.709, p < 0.0001, η2 = 0.817; E4: n = 145, F_1.880, 270.737_ = 771.225, p < 0.0001, η2 = 0.843; E6: n = 67, F_1.618, 106.807_ = 636.019, p < 0.0001, η2 = 0.906; BZ: n = 95, F_1.602, 150.625_ = 535.607, p < 0.0001, η2 = 0.851; 2H: n = 81, F_1.786, 142.899_ = 518.47, p < 0.0001, η2 = 0.866). Post hoc analyses confirm responses are significantly lower at each time point when compared to the preceding session across all odors (Figure 3B) as they are for the Odor only group. This implies additional exposure to E5 during the Odor only treatment or exposure to shock alone does not disproportionately affect some odors.

**Figure 3.**
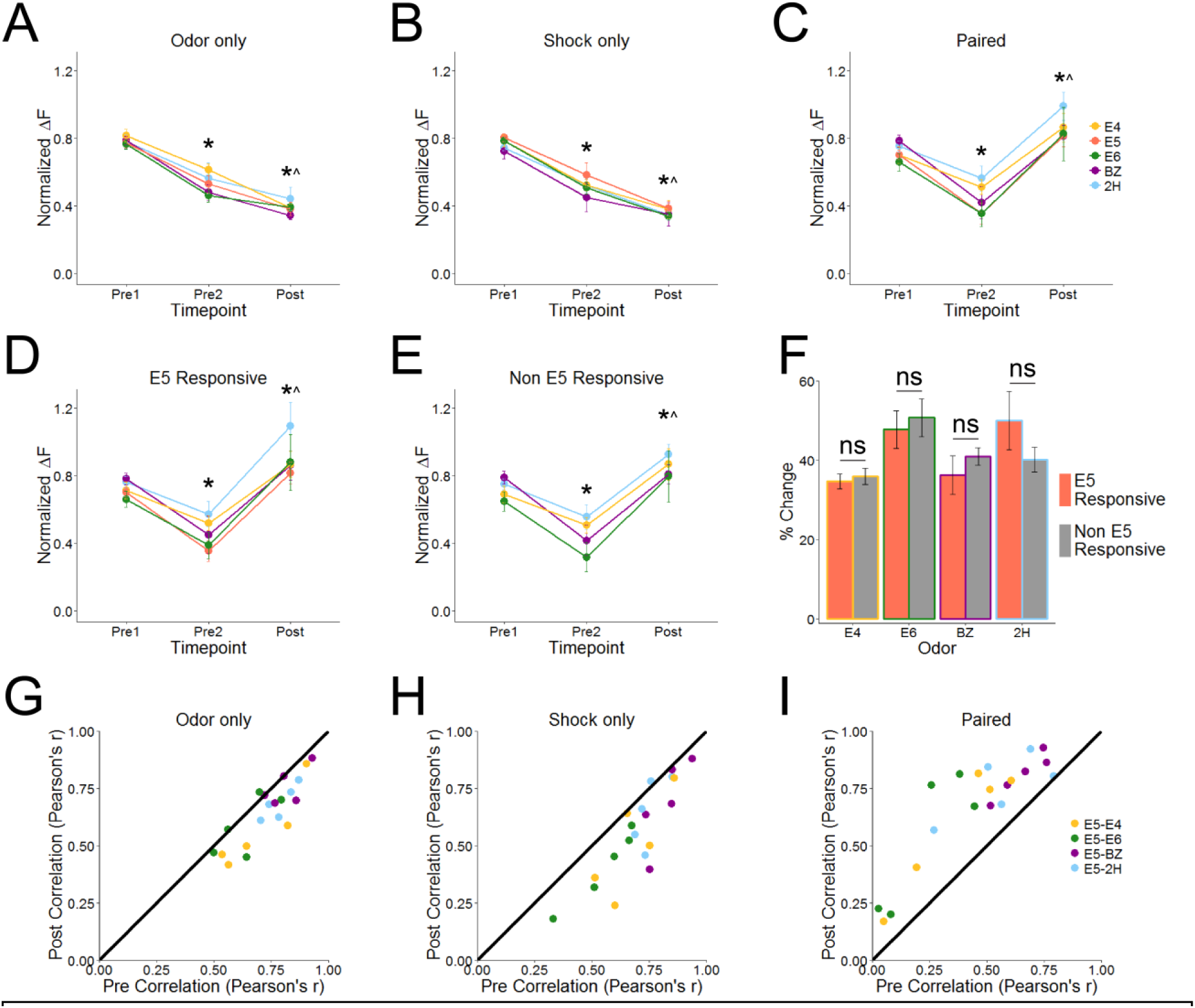
Enhanced responses are global, not odor- or glomerulus-specific. (A-C) Normalized glomerular responses over time for each imaged odor. Responses for all odors continually decrease over time for Odor only (A) and Shock only mice (B). Glomerular responses to all odors of Paired mice (C) decrease before learning (from Pre1 to Pre2) followed by robust reinstatement of responses after learning (Post). The same learning-induced enhancement in Paired mice occurs in E5 Responsive (D) and Non E5 Responsive (E) glomeruli, indicating glomerular overlap does not play a significant role in learning-induced alterations (F). (G-H) Scatterplots showing the average correlation of spatial glomerular activation patterns between E5 and each of the neutral odors before (Pre2, x-axis) and after (Post, y-axis) for all Odor only (G), Shock only (H), and Paired (I) mice. Solid black line represents theoretical “no change” line. Activation patterns of E5 and neutral odors decorrelate after training in both Odor only (G) and Shock only (H) mice but become more correlated after training in Paired mice (I). Data presented mean ± sem. *p < 0.001 from Pre1, ^p < 0.001 from Pre2

When exploring glomerular responses for different odors in the Paired group, we find all odors display the same pattern of decreased responses from Pre1 to Pre2 followed by robust potentiation at the Post time point (RM ANOVA E5: n = 149, F_1.767, 261.512_ = 435.719, p < 0.0001, η2 = 0.746; E4: n = 154, F_1.601, 244.962_ = 330.582, p < 0.0001, η2 0.684; E6: n = 82, F_1.389, 112.483_ = 107.171, p < 0.0001, η2 = 0.570; BZ: n = 106, F_2, 210_ = 208.243, p < 0.0001, η2 = 0.665; 2H: n = 95, F_1.616, 151.858_ = 90.222, p < 0.0001, η2 = 0.490). Post hoc analyses confirm responses at Pre2 are lower than those at Pre1 for all odors and Post responses are significantly higher than those at both Pre2 for all odors (Figure 3C). In addition, Post responses are significantly elevated above those on Pre1 for all odors except BZ, where Post and Pre1 responses are statistically equal (p = 0.346). This demonstrates that the associative odor-foot shock learning, rather than just shock or odor experience alone non-specifically alters glomerular responses to odors.

Many glomeruli responsive to non-conditioned odors (E4, E6, BZ, and 2H) were also responsive to presentations of E5; therefore, we additionally investigated the extent to which glomerular overlap plays a role in the response changes described above. We calculated the percent change from Pre2 to Post for each glomerulus and then characterized them as “E5 Responsive” or “Non E5 Responsive”, which allowed us to directly compare how much responses change based on whether glomeruli overlap with E5. Responses of E5 glomeruli for Odor only and Shock only groups decreased 15.230% ± 1.019% and 21.350% ± 1.281% from Pre2 to Post, respectively, while E5 glomeruli in the Paired group increased 46.273% ± 1.636% from Pre2 to Post. In the Odor only group, there was no effect of overlap on percent change from Pre2 to Post for any odor (Independent samples t test, E4: t_134_ = 0.279, p = 0.781; E6: t_50.286_ = −1.000, p = 0.322; BZ: t_45.139_ = 0.838, p = 0.407; 2H: t_99_ = 0.422, p = 0.674). In the Shock only group, there was also no effect of overlap on percent change from Pre2 to Post for any odor except BZ (Independent samples t test, E4: t_140.224_ = −0.532, p = 0.595; E6: t_65_ = 0.265, p = 0.792; BZ: t_93_ = −2.662, p = 0.009; 2H: t_79_ = 0.007, p = 0.995). Finally, there was no significant difference in the Pre2 to Post enhancement for glomerular responses to any odors in the Paired group (Independent samples t test, E4: t_152_ = −0.457, p = 0.0648; E6: t_80_ = −0.434, p = 0.666; BZ: t_104_ = −0.943, p = 0.348; 2H: t_93_ = 1.358, p = 0.178), signifying that the response properties of individual glomeruli responsive to E5 were not altered during training (Figure 3F), thereby causing the non-specific enhancement described above. Rather, olfactory fear learning induces a global enhancement of all glomeruli, independent of odorant and overlap with the CS.

We next investigated whether the changes in individual glomerular responses following training altered the overall representation of non-conditioned odors (E4, E6, BZ, and 2H) to be more or less similar to the conditioned odor (E5). To accomplish this, we generated the averaged glomerular response maps of each odors and asked how well correlated the spatial pattern of activation was between each of the non-conditioned odors and E5 before and after training. Our analysis suggests that non-conditioned odors are equally as correlated or decorrelated with E5 after training for both Odor only and Shock only mice (Figures 3G and 3H). In contrast, the pattern of spatial activity in response to non-conditioned odors is more similar to the response elicited by presentations of the CS after training for Paired mice than it was before.

### Olfactory fear learning induces long-lasting behavioral fear and enhanced glomerular responses

In order to evaluate the duration of the post-learning effects, we repeated the Paired condition of the previous experiment but tested and imaged 72, rather than 24, hours after foot-shock training (Figure 4A). There was a main effect of stimulus on behavioral freezing following fear learning (ANOVA: F_5, 18_ = 6.677, p = 0.001, η2 = 0.650). Post hoc analysis indicates only baseline freezing was significantly different from freezing to the CS (E5), p < 0.0009, indicating the odor-shock training paradigm produces broad fear generalization across all odors up to 72 hours after training takes place (Figure 4B).

**Figure 4.**
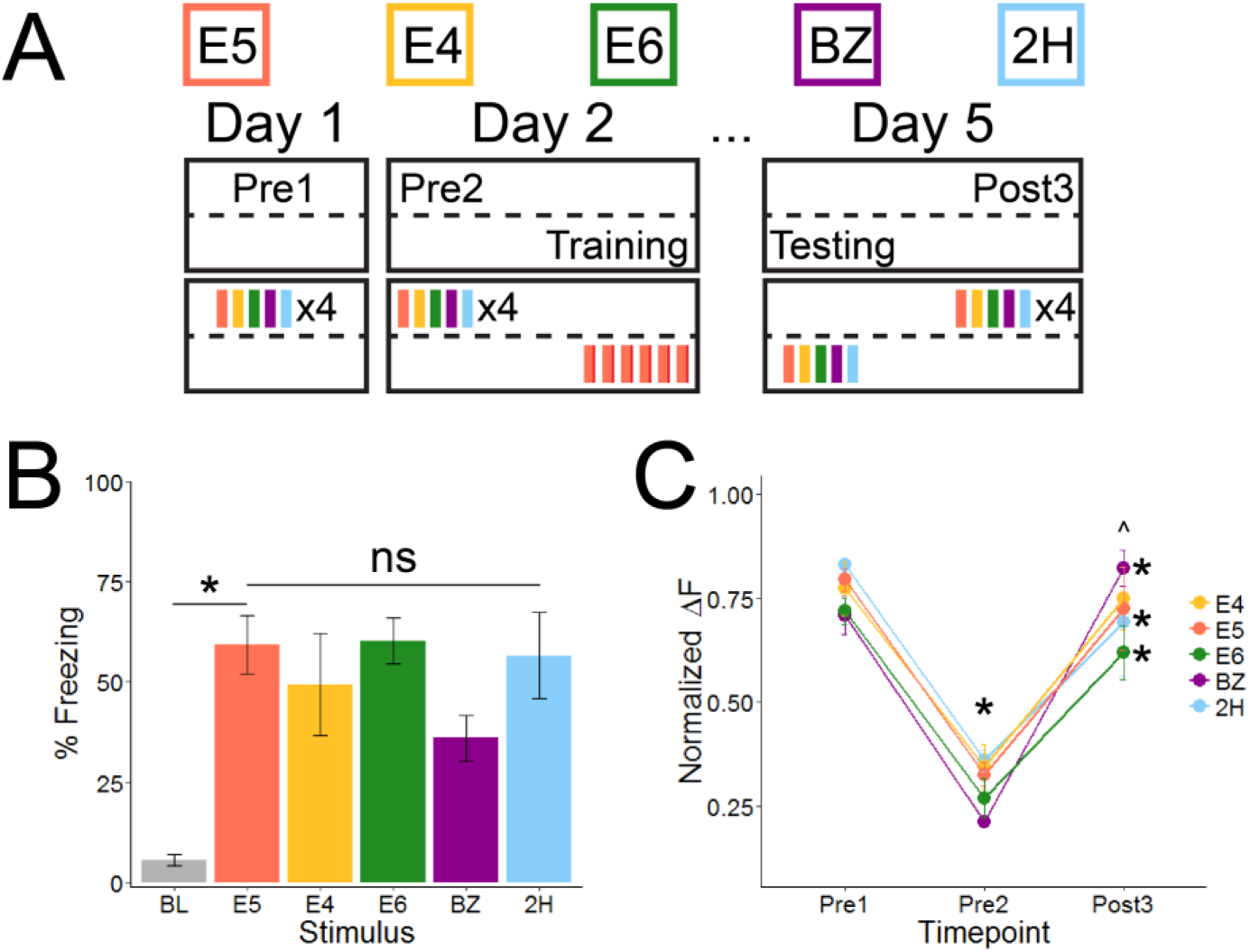
Olfactory fear learning induces long-lasting behavioral fear and glomerular enhancements. (A) Timeline detailing the odors used (top) and both imaging (above dotted line) and behavioral (below dotted line) experiments. (B) 72 hours after training, mice were exposed to each of the 5 odors and freezing was measured. Mice froze significantly more to E5 than baseline, indicating acquired fear to the CS. Mice generalized fear across all other tested odors (freezing to other odors not significantly different from freezing to E5). (C) Glomerular responses to all odors significantly decrease from Pre1 to Pre2; however, even 72 hours after training, responses are significantly greater after learning (Post) than before (Pre2). Data presented mean ±sem. *p < 0.001 from Pre1, ^p < 0.001 from Pre2

The same mice also underwent awake imaging 72 hours after training (Post3) to characterize whether learning-induced glomerular enhancements were long lasting. There was a main effect of time on global glomerular responses (RM ANOVA: n = 401 averaged glomerular responses, F_1.529, 611.410_ = 941.730, p < 0.0001, η^2^ = 0.702), in which these mice displayed the characteristic response decrease from Pre1 and Pre2 and an enhancement of glomerular responses following training that was still visible 72h later (Post3). This pattern is not driven by any particular odor (RM ANOVA E5: n = 107, F_1.642, 174.104_ = 230.484, p < 0.0001, η^2^ = 0.685; E4: F_1.542, 161.995_ = 222.538, p < 0.0001, η^2^ = 0.679; E6: F_1.547, 94.374_ = 202.332, p < 0.0001, η^2^ = 0.768; BZ: F_1.705, 78.445_ = 284.862, p < 0.0001, η^2^ = 0.861; 2H: F_1.310, 102.161_ = 225.059, p < 0.0001, η^2^ = 0.743). In fact, post hoc analyses indicate that responses to each odor on Post3 were significantly higher than responses for the same odor on Pre2 (Figure 4C; p < 0.001). The sustained, much like the initial, enhancement, is not odor specific, indicating that olfactory fear learning globally increases glomerular responses in a long-lasting manner.

### Anesthetized mice display weaker glomerular enhancements and suppression following olfactory fear learning

We next examined whether wakefulness modulates the olfactory learning-induced glomerular response profile. We repeated the Paired condition in the first set of experiments but completed each of the imaging sessions in anesthetized, rather than awake, behaving mice. We confirmed olfactory fear learning in these mice by testing their awake behavioral freezing to each of the odors 24 hours following odor-shock conditioning (Figure 5A; ANOVA: F_5, 18_ = 3.224, p = 0.030, η2 = 0.472). Post hoc analyses demonstrates that only freezing during the baseline minute is significantly different from freezing to E5 (p = 0.007), suggesting broad behavioral generalization is retained even in the anesthetized condition.

**Figure 5.**
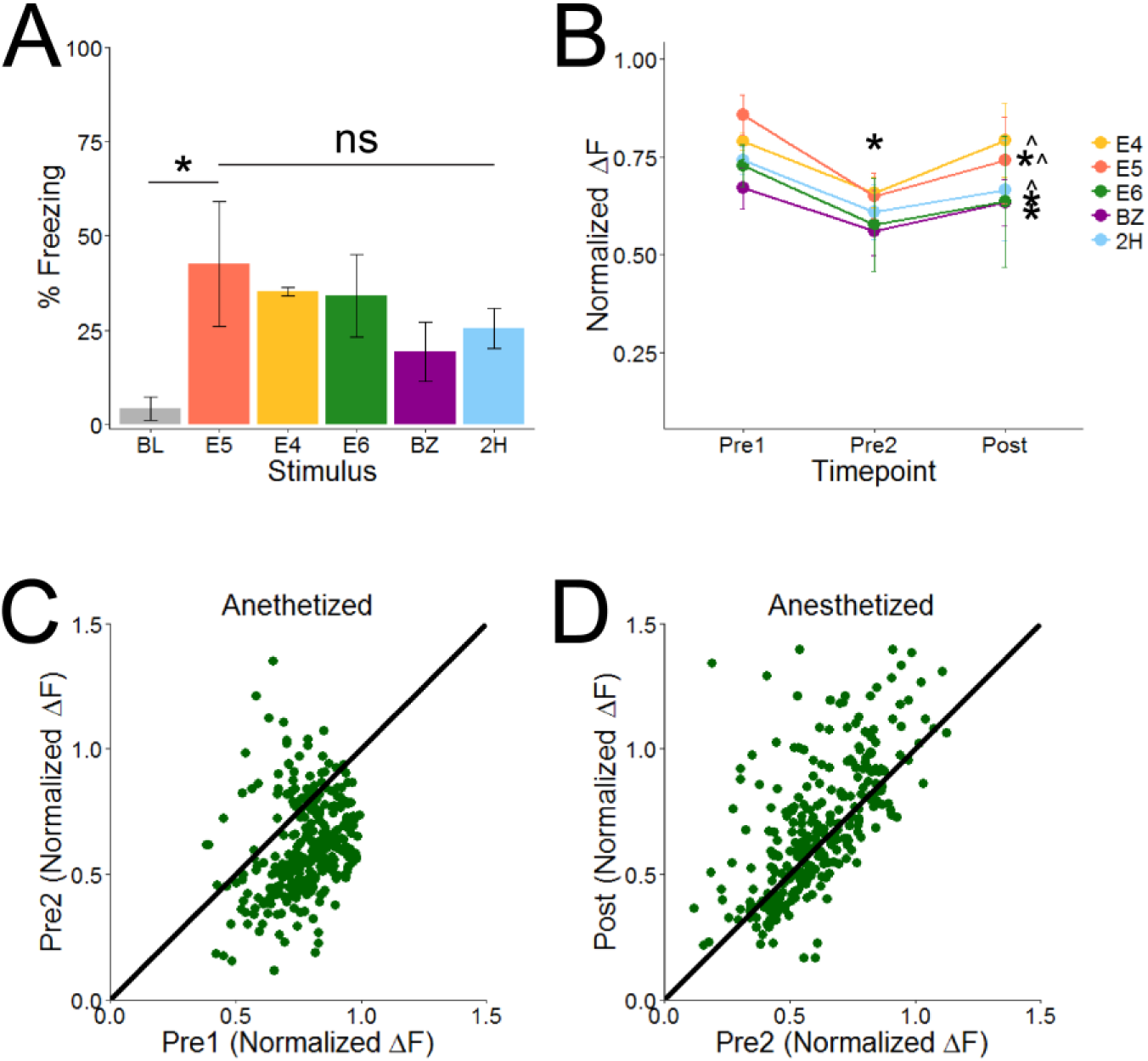
Learning-induced glomerular changes are variable in anesthetized mice. (A) 24 hours after training, mice froze significantly more to E5 than baseline, indicating acquired fear and generalized fear across all other tested odors (freezing to other odors not significantly different from freezing to E%). (B) Glomerular responses of anesthetized mice decreased from Pre1 to Pre2. After training (Post), only averaged responses of E5, E4, and 2H were significantly enhanced (relative to Pre2), while responses to E6 and BZ were not significantly different. (C&D) Scatterplots showing normalized Pre1 (x-axis)/Pre2 (y-axis) responses (C) or Pre2 (x-axis)/Post (y-axis) responses (D) for each recorded glomerulus from Anesthetized mice. Solid black line represents theoretical “no change” line. Glomerular responses generally decrease from Pre1 to Pre2, as evidenced by the majority of the points falling below the no change line (C). On average, responses slightly increase from Pre2 to post; however, of the 292 glomeruli analyzed in the anesthetized mice, only 59.9% were enhanced after training while 40.1% were suppressed. Data presented mean ± sem. *p < 0.001 from Pre1, ^p < 0.001 from Pre2

Mice in the anesthetized condition also display decreased responses from Pre1 to Pre2 (Figures 5B and 5C) followed by learning-induced glomerular enhancements (RM ANOVA: n = 292 averaged glomerular responses, F_1.496, 435.357_ = 68.702, p < 0.0001, η^2^ = 0.191); however, we noted the enhancement appeared reduced compared to the awake Paired group. A scatterplot demonstrated suppression of several glomeruli after training (Figure 5D), which occurred in all odors (RM ANOVA E5: n = 75, F_1.492, 110.435_ = 24.124, p < 0.0001, η2 = 0.246; E4: n = 80, F_1.315, 103.906_ = 22.307, p < 0.0001, η2 = 0.220; E6: n = 58, F_1.297, 73.934_ = 11.469, p < 0.0005, η^2^ = 0.168; BZ: n = 24, F_2, 46_ = 6.241, p < 0.004, η2 = 0.213; 2H: n = 55, F_1.409, 76.098_ = 12.232, p < 0.0002, η2 = 0.185). In addition, post hoc analyses revealed glomerular responses to both E6 and BZ were not significantly different from Pre2 to Post (p = 0.135 and 0.231, respectively). Out of the 292 glomeruli analyzed in the anesthetized mice, only 59.9% (175 total; E5 = 47, E4 = 56, E6 = 34, BZ = 13, 2H = 25) were enhanced after training and 40.1% (117 total; E5 = 28, E4 = 24, E6 = 24, BZ = 11, 2H = 30) were decreased. Comparatively, in the awake Paired condition only 2 of the 586 glomeruli (0.34%) analyzed exhibited lower responses after training. Also of note, the decreased responsivity from Pre1 to Pre2 is qualitatively smaller in anesthetized than awake mice. This illustrates wakefulness modulates both passive experience- and learning-induced glomerular alterations and that anesthetized imaging in the context of experience-dependent glomerular transformations do not mirror that seen in the awake condition.

### Post-training glomerular response enhancements are independent of general fear states

One possible explanation for post-training glomerular response alterations in awake mice that differ from those observed in anesthetized mice is that the imaging paradigm requires continually presenting fear-inducing stimuli to the mice in order to assess post-training changes. It is possible that this repeated exposure induces a general fear state that enhances glomerular responses. We devised a paradigm to image glomerular responses to odors in the presence and absence of a fear inducing stimulus (Figure 6A). Instead of fear conditioning Paired mice to an odor, we conditioned them to a 10kHz, 82dB tone. Additionally, we split the Post time point into two halves (Post1 and Post2). In the first half (Post1), we imaged odor responses normally; however, in the second half (Post2), we imaged odor responses immediately after a 10s presentation of the conditioned tone.

**Figure 6.**
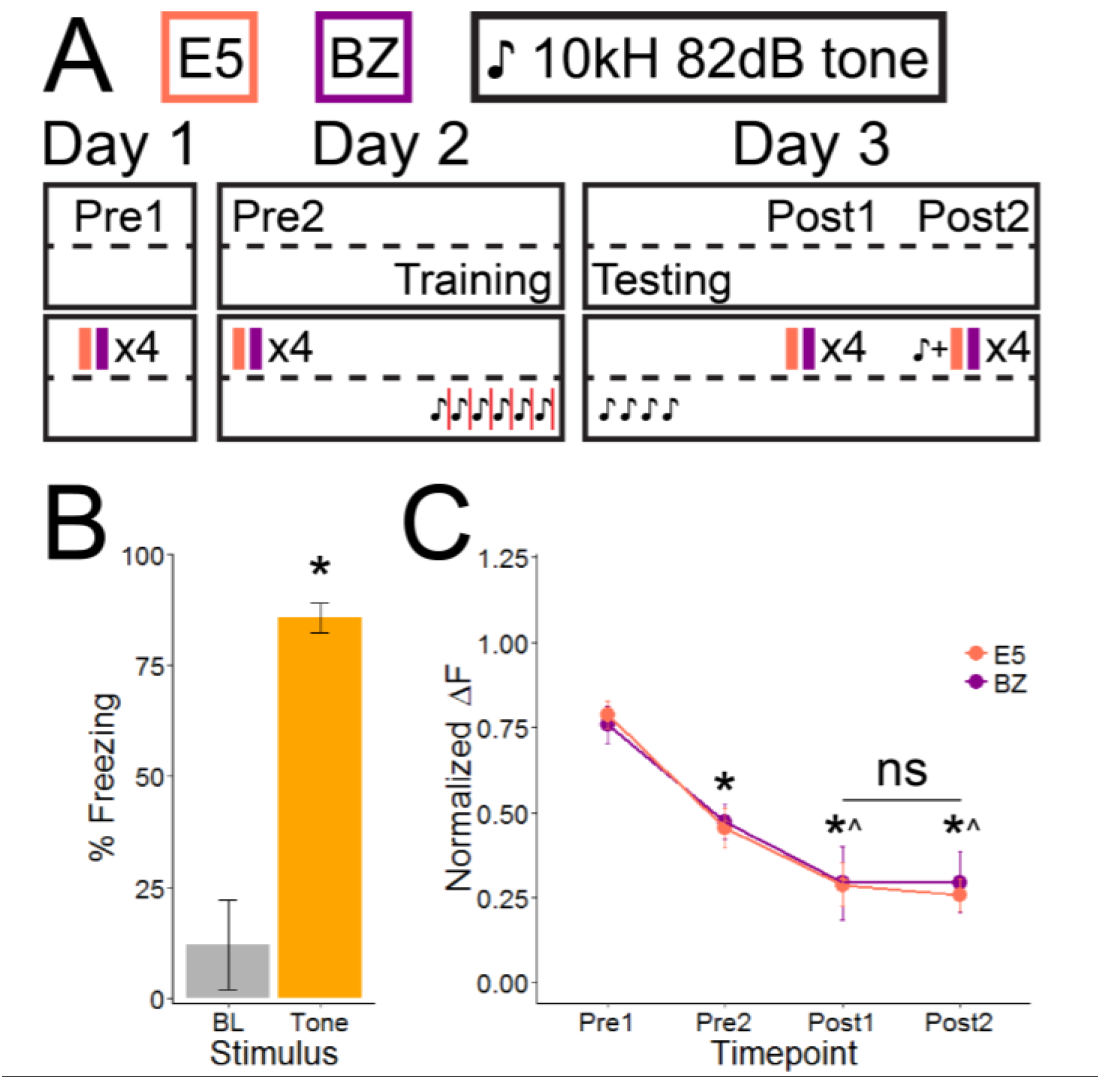
The expression of learning-induced glomerular response enhancements is independent of general fear states. (A) Timeline detailing the stimuli used (top) and both imaging (above dotted line) and behavioral (below dotted line) experiments. (B) Tone-shock conditioned mice learn to fear the conditioned tone and freeze to it significantly more than baseline. (C) Odor-evoked glomerular responses of mice conditioned to fear a tone decreased from Pre1 to Pre2 and again from Pre2 to Post1. Importantly, there was no significant difference between odor-evoked glomerular responses during Pre1, when awake mice were imaged normally, and Post2, when we experimentally induced fear by preceding each odor imaging trial with a presentation of the fear-inducing tone. Data presented mean ± sem. p<0.001 from Pre1, ^p < 0.001 from Pre2

Tone-shock conditioned produced robust freezing to presentations of the conditioned tone relative to baseline freezing in the first minute (Figure 6B; Independent samples t test: n = 3, t_4_ = −6.650, p < 0.003). Glomerular responses significantly decreased across imaging sessions (Figure 6C; RM ANOVA: n = 128 averaged glomerular responses, F_2.317, 294.304_ = 729.000, p < 0.0001, η2 = 0.852) and the decrease was consistent across the two tested odors (RM ANOVA E5: n = 72, F_2.215, 157.276_ = 544.812, p < 0.0001, η2 = 0.885; BZ: n = 56, F_1.792, 98.583_ = 239.018, p < 0.0001η^2^= 0.812). Post hoc analyses indicate all time points are significantly different from one another except Post1 and Post2 (p = 1.000, for both odors). The lack of significant difference between recorded responses at Post1 (in absence of fear inducing tone) and Post2 (in presence of fear inducing tone) suggest a global fear state is not responsible for the augmented responses observed in odor-shock conditioned mice, but that the enhancement is likely due to fear learning itself.

### Global, but not CS-specific, glomerular enhancements are fear learning dependent

Because the post-training enhancements could not be attributed to a global fear state that might simply strengthen all incoming sensory information, we next evaluated whether glomerular enhancements were dependent upon olfactory fear learning. We predicted that infusing muscimol (MUSC) into the basolateral amygdala (BLA) prior to odor-shock training would inactivate the BLA, thus interrupting fear learning (Figure 7A). Mice that received infusions of vehicle immediately before training demonstrate typical fear learning (Figure 7B; ANOVA: n = 5, F_5, 24_ = 8.703, p < 0.0001, η^2^ = 0.645). Post hoc analyses reveal mice freeze significantly more to E5 than baseline (p < 0.001) and generalize fear from E5 to E4, E6, and 2H, but not to BZ (p = 0.020, all others not significantly different from E5). On the other hand, MUSC mice did not exhibit signs of fear learning (Figure 7C; ANOVA: n = 5, F_5, 24_ = 1.107, p = 0.383, η2 = 0.187), confirming our hypothesis that infusions of MUSC into the BLA before training could effectively block fear learning.

**Figure 7.**
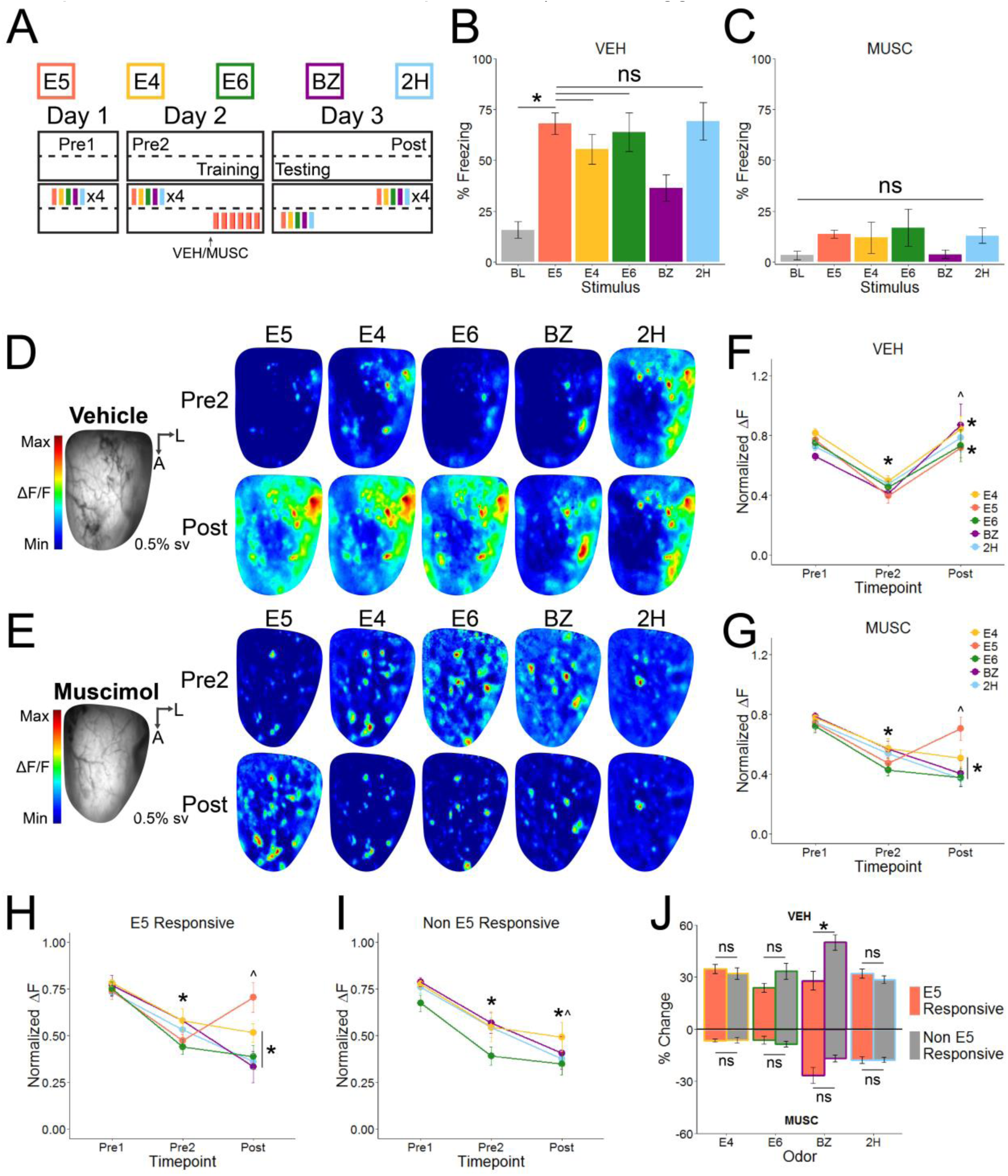
Generalized, but not CS-specific, glomerular enhancements are associative learning-dependent. (A) Timeline detailing the odors used (top), timing of drug infusion, and both imaging (above dotted line) and behavioral (below dotted line) experiments. (B&C) Mice were exposed to all 5 odors 24 hours after training and freezing was measured. In mice receiving VEH infusions before training, presentations of E5 elicited significantly more freezing than baseline, indicating they learned to fear the CS. Additionally, VEH mice generalized fear to all odors, except BZ. In contrast, mice receiving MUSC infusions before training, did not freeze significantly more to presentations of E5 than baseline, indicating they did not learn. (D and E) RLI frames and psuedocolored averaged Pre2 and Post maps from representative VEH (D) and MUSC (E) mice. (F&G) Normalized glomerular responses over time for each imaged odor for all VEH (F) and MUSC (E) mice. Responses for all odors in both groups significantly decrease from Pre1 to Pre2. Responses to all odors are significantly increased after training in VEH mice. In contrast, only responses to presentations of E5 are enhanced in MUSC mice; all other odor responses are continually suppressed. In MUSC mice, the same post-training suppression of non-conditioned odors occurs regardless of whether glomeruli are E5 Responsive (H) or Non E5 Responsive (I). (J) Glomerular overlap does not affect the percent change of glomerular responses (Pre2 to Post) in VEH (top; with the exception of BZ) or MUSC (bottom) mice, indicating glomerular changes are odor-rather than glomerulus-specific (J). Data presented mean ± sem. *p < 0.001 from Pre1, ^p < 0.001 from Pre2

Not surprisingly, glomerular responses of VEH mice differed over the imaging sessions (RM ANOVA: n = 596 averaged glomerular pairs, F_1.384, 823.392_ = 754.435, p < 0.0001, η^2^ = 0.559), with post hoc analyses confirming a significant decrease in responses from Pre1 to Pre2 and a significant increase from Pre2 to Post (Figures 7D), similar to that of non-cannulated Paired mice. In general, glomerular responses of MUSC mice changed over time (RM ANOVA: n = 612 averaged glomerular responses, F_1.767, 1079.568_ = 599.623, p < 0.0001, η2 = 0.495) and post hoc analyses revealed that responses decreased from Pre1 to Pre2 with an additional, statistically significant decrease from Pre2 to Post (Figure 7E; p < 0.001). The lack of post-training enhancement in the MUSC group suggests these changes are learning-dependent; however, we noticed not all glomerular responses in the MUSC group were further decreased following odor-shock training. To determine whether responses to individual odors were differentially affected after training with infusions, we analyzed the responses of each odor individually.

In the VEH group, all odors presented similar patterns of responsivity over the imaging sessions (RM ANOVA E5: n = 140, F_1.628, 226.249_ = 393.528, p < 0.0001, η2 = 0.739; E4: n = 141, F_1.335, 186.900_ = 190.665, p < 0.0001, η^2^ = 0.577; E6: n = 91, F_1.369, 123.206_ = 119.941, p < 0.0001, η^2^ = 0.571; BZ: n = 99, F_1.206, 118.164_ = 98.089, p < 0.0001, η2 = 0.500; 2H: n = 125, F_1.654, 205.090_ = 209.034, p < 0.0001, η^2^= 0.628) with post hoc analyses verifying that Pre2 responses are significantly lower than Pre1, and Post responses are significantly higher than Pre2 for all 5 odors (Figure 7F; p < 0.001). In contrast, not all odor responses in MUSC mice continue to decrease after training (RM ANOVA E5: n = 160, F_1.622, 257.85_ = 195.242, p < 0.0001, η2 = 0.551; E4: n = 156, F_1.543, 239.236_ = 284.548, p < 0.0001, η2 = 0.647; E6: n = 89, F_1.788, 157.361_ = 234.108, p < 0.0001, η2 = 0.727; BZ: n = 85, F_2, 168_ = 179.119, p < 0.0001, η2 = 0.681; 2H: n = 122, F_2, 242_ = 579.7, p < 0.0001, η2 = 0.827). Post hoc analyses reveal responses to all odors decrease from Pre1 to Pre2 (p < 0.001 for all five odors) but only glomerular responses to non-conditioned odors (E4, E6, BZ, and 2H) display further suppression from Pre2 to Post (Figure 7G; p < 0.001), while glomerular responses to the conditioned odor, E5, are enhanced after odor-shock training (p < 0.001). In fact, Post responses to E5 are not significantly different from those measured on Pre1 (p = 0.14), indicating full reinstatement of the initial E5 response after fear conditioning even in the absence of fear learning. These results illustrate that generalized, or global, glomerular enhancement is learning-dependent, while CS-specific enhancements do not require learning.

Many glomeruli responsive to E5 are also responsive to presentations of the other, non-conditioned odors. Further, responses to E5 were the only ones enhanced after training in MUSC mice. Therefore, we wanted to explore whether all “E5 Responsive” glomeruli might be enhanced, but the effect obscured by averaging the responses of “E5 Responsive” with “Non E5 Responsive” glomeruli. We calculated percent change from Pre2 to Post for each glomerulus and then classified them based on whether they responded to presentations of the CS, which allowed us to directly compare the effects of E5 overlap. When mice received VEH infusions prior to odor-shock training, E5 glomerular responses increased 31.364% ± 1.500% from Pre2 to Post. Glomerular overlap with E5 appeared to have no effect on the amount of observed post-training change for most of the non-conditioned odors (Figure 7H; Independent samples t test E4: t_139_ = 0.591, p = 0.555; E6: t_89_ = −1.895, p = 0.061; BZ: t_97_ = −2.428, p = 0.017; 2H: t_123_ = 1.046, p = 0.298) with only BZ exhibiting a significant difference between “E5 Responsive” and “Non E5 Responsive” in terms of percent change. E5 glomeruli of MUSC mice increased their responses 23.139% ± 1.492% after training, while responses to all other odors decreased. Glomerular overlap with E5 did not impact the change from Pre2 to Post for any of the non-conditioned odors (Figure 7I; Independent samples t test E4: t_154_ = −0.153, p = 0.879; E6: t_85.762_ = 0.797, p = 0.428; BZ: t_83_ = - 1.537, p = 0.128; 2H: t_120_ = −0.047, p = 0.963). This suggests a mechanism for odor rather than glomerulus-specific modulation in the OB following odor-shock training, even in the absence of fear learning (Figure 7J). Taken all together, the results of this experiment provide evidence for two separate mechanisms that modulate OB glomerular responses. One is specific to the CS and does not require learning to regulate responses, while the second is learning-dependent and results in a global, or non-specific, gain across OB glomeruli.

### Suppressing fear centers during expression does not suppress learning-induced glomerular enhancements

Finally, we explored whether the post-training glomerular enhancements could simply be due to amygdalar activation as a result of a fear state that increased sensory information in a modality-specific manner. To directly compare post-learning glomerular responses, we split the Post time point into two halves (Post1 and Post2). In the first half (Post1) we imaged odor responses normally; however, 10 minutes before the second half (Post2), we infused either VEH or MUSC into the BLA and imaged odor responses while the BLA was normally active or inactivated, respectively (Figure 8A). Both groups of mice were subjected to odor-shock training, resulting in significant behavioral freezing (Figures 8B and 8C; ANOVA VEH: n = 5, F_5, 24_ = 9.056, p < 0.0001,η^2^ = 0.654; ANOVA MUSC: n = 5, F_5, 24_ = 17.437, p < 0.0001, η^2^ = 0.784). Post hoc analyses establish significantly higher freezing to E5 than baseline (p < 0.001, both groups), with broad generalization from E5 to all other odors in VEH mice (p > 0.065) and generalization from E5 to all odors except BZ (p < 0.001) in MUSC mice. These behavioral results are similar to the VEH results obtained in the previous experiment, which also contained mice that had bilateral cannula implantation in the BLA.

**Figure 8.**
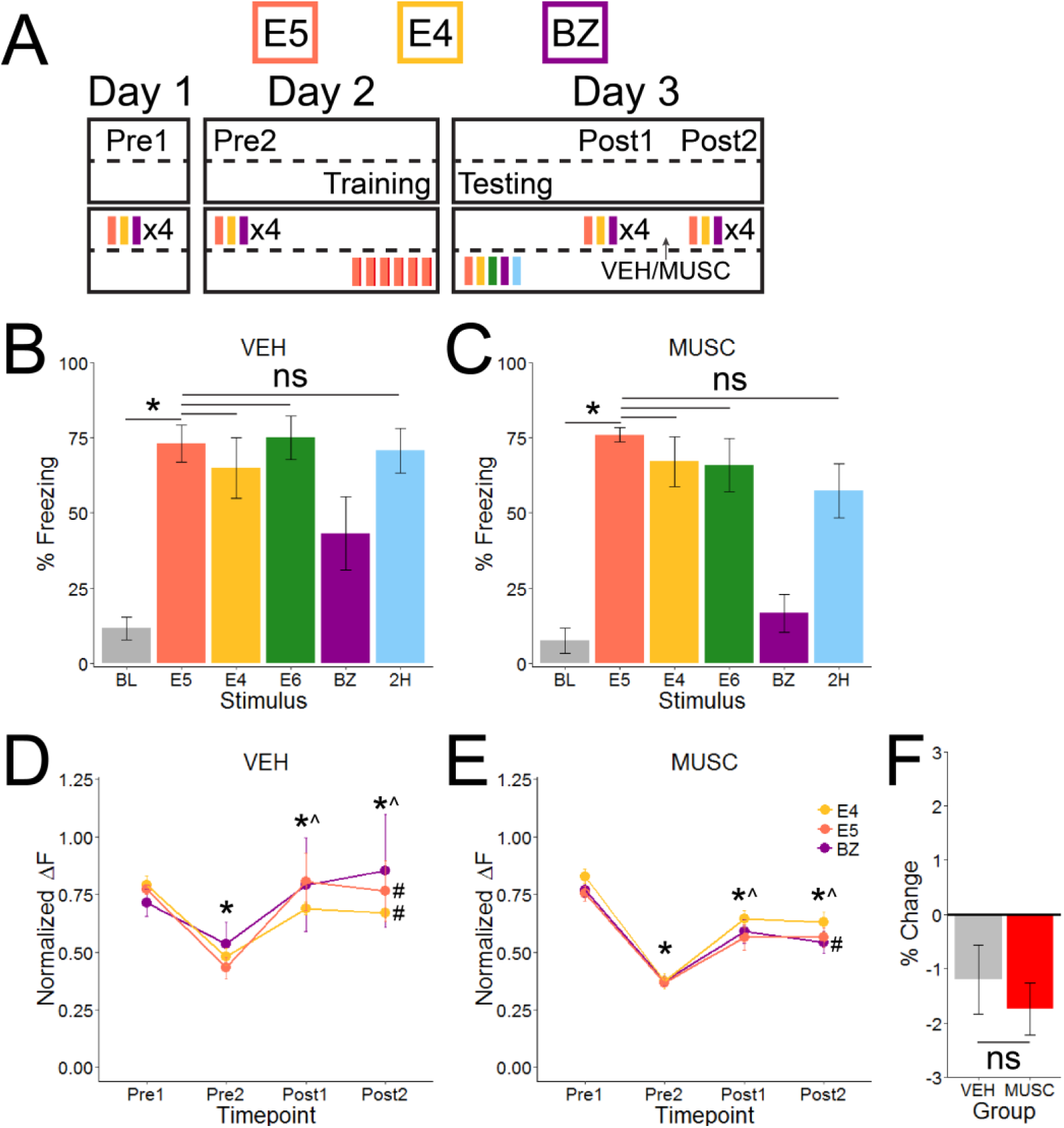
Amygdala inactivation during expression of learning does not impact glomerular responses. (A) Timeline detailing the odors used (top), timing of drug infusion, and both imaging (above dotted line) and behavioral (below dotted line) experiments. (B&C) Freezing to all 5 odors was measured 24 hours after training and confirms that both groups of mice froze significantly more to the CS than baseline and generalized that fear to all odors except BZ. (D&E) Averaged glomerular responses for both groups decreased significantly from Pre1 and Pre2, but were reinstated after learning (Post1). Infusions of either VEH or MUSC occurred between Post1 and Post2. Following infusions, both groups exhibited mixed profile responses, with some odor responses increasing, others decreasing, and some not changing significantly. Additional analysis quantifying the average % change from Post1 to Post2 (F) indicates no significant difference between groups, indicating BLA inactivation does not alter glomerular responses in a meaningful way. *p < 0.001 from Pre1, ^p < 0.001 from Pre2, #p <0.001 from Post1

Glomerular responses of VEH mice changed over time (RM ANOVA: n = 367 averaged glomerular responses, F_1.477, 540.494_ = 224.211, p < 0.0001, η^2^ = 0.380), with post hoc analyses demonstrating a significant decrease in responses from Pre1 to Pre2 (p < 0.001), rebound of glomerular responses from Pre2 to Post1 (p < 0.001), and a small decrease from Post1 to Post2, after VEH infusion into the BLA. Glomerular responses of MUSC mice changed similarly over the imaging sessions (RM ANOVA: n = 379 averaged glomerular responses, F_2.151, 813.016_ = 879.834, p < 0.0001, η^2^ = 0.700), and post hoc analyses reveal a significant suppression of responses from Pre1 to Post2 (p < 0.001), enhancement from Pre2 to Post1 (p < 0.001) and a significant decrease from Post1 to Post2 (p = 0.002) after MUSC infusion. We noticed that all odors for both groups follow the same trend of decreasing from Pre1 to Pre2, with a rebound of the response from Pre2 to Post1, but that individual odors exhibit mixed response alterations between Post1 and Post2. Therefore, we analyzed each odor separately over time (RM ANOVA VEH E5: n = 139, F_1.359, 187.490_ = 144.905, p < 0.0001, η^2^ = 0.512; E4: n = 142, F_2.212, 311.887_ = 252.805, p < 0.0001, η^2^ 0.642; BZ: n = 86, F_1.291, 109.740_ = 19.612, p < 0.0001, η^2^ = 0.187; RM ANOVA MUSC E5: n = 145, F_1.980, 285.160_ = 307.731, p < 0.0001, η^2^ = 0.681; E4: n = 145, F_2.478, 356.836_ = 494.151, p < 0.0001, η^2^ = 0.774; BZ: n = 89, F_1.689, 148.633_ = 145.011, p < 0.0001, η^2^ = 0.622). In doing so, we discovered that responses to E5 and E4 were significantly decreased (p < 0.001 and 0.02, respectively), but BZ responses were not significantly different (p = 0.116) from Post1 to Post2 in mice receiving VEH infusions (Figure 8D). In mice receiving MUSC infusions, responses to E5 and E4 were not significantly different (p = 1.000 and 0.126, respectively), but BZ responses were significantly decreased (p < 0.001) from Post1 to Post2 (Figure 8E).

In light of the mixed response profiles from Post1 to Post2 in both groups, we decided to test whether the response changes between these imaging session halves was dependent upon experimental condition (infusion of either VEH or MUSC between the two halves). To do so, we calculated the percent change for each glomerulus from Post1 to Post2. The response decrease from Post1 to Post2 was not significantly different between groups (Independent samples t test: t_744_ = 0.679, p = 0.497, Mean ± SEM: VEH = −1.202% ± 0.634%, MUSC = −1.739% ± 0.480%), indicating inactivation of the BLA after learning did not reduce glomerular responses relative to VEH control responses (Figure 8F).

## Discussion

Pairing awake*in vivo* calcium imaging with behavior, we investigated the effect of olfactory aversive learning on glomerular odor responses, including their spatial coding. The results demonstrate that odor familiarity leads to reduced glomerular responses that are evident as early as the second day of odor exposure. These responses continue to decrease over days, with olfactory fear learning significantly and long-lastingly enhancing glomerular layer odor representations for the CS as well as neutral, non-conditioned odors. Enhancements are not specific to CS-responsive glomeruli, indicating non-specific potentiation across glomeruli. Moreover, the spatial representation of non-conditioned odors becomes more correlated to the CS representation, possibly contributing to broad behavioral fear generalization by increasing perceptual similarity. Increased glomerular responses following fear conditioning are not caused by altered respiratory responses or global fear states as a result of shock learning, nor can it be suppressed by inactivating the BLA after learning takes place. Chiefly, this study demonstrates two distinct mechanisms responsible for glomerular enhancements: an associative learning-independent mechanism, which supports CS-specific enhancements, and an associative learning-dependent mechanism, which promotes non-specific potentiation associated with generalization. Overall, this study demonstrates that sensory representations of learned stimuli are altered at the earliest stages of sensory processing. These sensory processing changes are potentially large enough to alter stimulus perception and could constitute a basis for behavioral outcomes of learning. To our knowledge, this is the first report of distinct mechanisms that mediate specific vs generalized stimulus response potentiation following classical fear learning.

These findings expand upon previous reports of learning induced changes in the OB and higher olfactory centers in insects (Blum et al., 2009; Chen et al., 2015; Faber et al., 1999), rodents (Barnes et al., 2011; Jones et al., 2008; Sevelinges et al., 2008), and humans (Li et al., 2008). Recent imaging studies in anesthetized rodents demonstrate increased responses to the trained odorant in both OSNs and post-synaptic M/T cells (Fletcher, 2012; Kass et al., 2013). Our anesthetized experiment confirms these findings, even with differences in experimental paradigms. In addition, we observed both increased and decreased post-training glomerular responses in our anesthetized experiment, as previously reported (Fletcher, 2012), while almost all glomeruli were significantly enhanced after training in awake mice. This likely reflects differences between anesthetized and awake conditions such as activity of intrabulbar inhibitory circuits or state-dependent centrifugal modulation (Blauvelt et al., 2013; Rothermel and Wachowiak, 2014); though it is also possible that robust expression of learning-induced glomerular plasticity requires wakefulness (Kato et al., 2012).

### Potential distinct mechanisms of experience-induced glomerular plasticity

We demonstrate, in the absence of any reinforcement, glomerular responses decrease across days. These findings are in line with previous studies establishing that stimulus familiarity leads to reductions in sensory responses (Buonviso and Chaput, 2000; Buonviso et al., 1998; Gdalyahu et al., 2012; Kato et al., 2012; McNamara et al., 2008). While we did not explore the cause of the non-associative learning suppression, previous work concludes this form of plasticity is generated by changes downstream of OSN input to glomeruli, as this effect is not seen when imaging OSN glomerular responses (Kato et al., 2012). The passive experience-dependent suppression is likely regulated by inhibitory interneurons within the OB that are reduced under anesthesia (Kato et al., 2012). Interestingly, we detect less suppression in anesthetized than awake mice, which is consistent with this idea.

We used MUSC to transiently inactivate the BLA during acquisition (Ribeiro et al., 2011; Wilensky et al., 1999), which prevented fear learning in mice receiving MUSC infusions but not those receiving infusions of VEH. Even though MUSC mice did not acquire learned fear of E5, they exhibited augmented glomerular responses to E5, indicating CS-specific potentiation does not require associative learning in this case. The fact that the CS-specific enhancements remain even when associative learning is blocked suggests that the first mechanism, which produces CS-specific enhancements, does not require associative learning. One possible explanation for this is experience-dependent structural changes within the glomerular layer itself. Fear conditioning can increase glomerular size (Jones et al., 2008); however, such structural changes are caused by increased number of OSN axons innervating glomeruli. We do not believe this drives our effect as the previous study trained mice over several days and weeks, allowing time for anatomical reorganization that our shorter training paradigm likely does not allow. Alternatively, CS-specific plasticity could occur in the glomerular layer downstream of OSNs. A previous report indicates odor-specific synaptic tagging in the OB glomerular and external plexiform layers, where dendrites and somas of M/T cells are located, as well as synaptic AMPA receptor insertion following odor experience (Modarresi et al., 2016), which could amplify glomerular responses. However, the enhancement we observed in the absence of learning was specific to E5, with no enhanced responses to other odors that activate those glomeruli. This suggests that the mechanisms responsible are not structural changes within OB neurons, as this should lead to enhanced responses to all odors that activate the glomerulus. Instead, the mechanism underlying the odor-specific enhancements likely involves changes within OB circuitry encoding E5. Odor exposure and learning can decrease activation of granule cells (Woo et al., 1996), which should result in disinhibition of M/T cells and increase activity within glomeruli (Johnson et al., 1995). Therefore, olfactory experience may modify the inhibitory circuits that tune M/T cell glomerular responses, which could shift responses in a highly odor-specific ensemble.

The second mechanism is a global, non-specific enhancement of all glomeruli that is dependent upon associative learning. Using an auditory fear paradigm in conjunction with OB imaging, we demonstrated that learning-induced enhancements are not a result of global fear states indiscriminately enhancing all incoming sensory information. However, it is possible that fear states only modulate sensory processing in a modality specific manner. Therefore, we additionally inactivated BLA by infusing MUSC to test whether suppressing fear centers during expression affects observed learning-induced glomerular enhancement. We detected no differences in glomerular responsivity to any of the tested odors between different BLA activity states. Together, these lines of evidence point to learning induced changes in centrifugal modulation of OB responsivity from higher brain regions as a likely candidate. There is a considerable amount feedback from cortical and neuromodulatory regions that can enhance OB responses to olfactory stimuli (Fletcher and Chen, 2010; Haberly and Price, 1978; Linster and Cleland, 2016; Mouret et al., 2009; Otazu et al., 2015; Price and Powell, 1970). Neuromodulatory systems are known to enhance representations of olfactory stimuli by acting on either local inhibitory interneurons or M/T cells. For example, both acetylcholine and serotonin release in the OB and reportedly enhance M/T cells odor responses (Bendahmane et al., 2016; Brunert et al., 2016; Kapoor et al., 2016; Rothermel et al., 2014). Both of these systems are involved in fear learning (Bauer, 2015; Wilson and Fadel, 2017) and could serve as the mechanism behind global increases in responsivity following fear conditioning.

### Potential impact

Previous literature suggests that changes in BLA coding correlates with generalization following aversive conditioning in non-human primates (Resnik and Paz, 2015); and studies in humans propose amygdala hyperactivity and exaggerated fear circuitry responses cause fear generalization (Morey et al., 2013; Shin et al., 2005). Here, we report that while BLA circuity is required for olfactory fear learning, BLA activity does not appear to affect glomerular responses after learning takes place, suggesting that the observed glomerular plasticity is not a result of amygdalar influence within the OB. While it is unlikely that fear generalization transpires as a direct cause of altered glomerular processing alone, such changes may play a crucial role in behavioral outcomes of learning.

Generalized learning-induced enhancement of glomerular responses could serve to increase perceptual similarity of experienced odors, as evidenced by increased representational correlations, and thereby contribute to behavioral fear generalization. Previous reports in the olfactory system demonstrate olfactory discrimination learning decorrelates odor responses in areas of olfactory cortex thought to code structural odor identity (Kadohisa and Wilson, 2006), possibly decreasing perceptual similarity and making fine discrimination between odorants easier. In fact, decorrelated cortical representations of odor mixtures predict behavioral discrimination of those mixtures (Barnes et al., 2011), reinforcing the idea that olfactory representational similarity confers perceptual similarity in a way that influences behavior. Similar learning-induced effects are reported in other sensory and model systems, confirming that sensory learning alters sensory processing and correlates with behavior (Edeline et al., 1993; Mukai et al., 2007; Mundy et al., 2014; Smith et al., 2015). Taken all together, any learning-induced transformation, even at the earliest stages of processing, that increases or decreases the representational similarity of sensory stimuli may prime generalized or specific behavioral responses, respectively.

The present results emphasize that fear learning increases perceptual similarity of sensory stimuli at the earliest stages of processing, which may bias downstream fear regions towards generalization. As disrupted fear generalization is a hallmark of anxiety and trauma and stressor-related disorders (Cahill and Foa, 2007; Lissek et al., 2014; Lissek et al., 2010), understanding the mechanisms and brain regions underlying fear generalization may inform future treatments of this pathological behavior as well as our basic understanding of the mechanisms underlying long-term memory.

## Acknowledgements

The authors want to thank Dr. Flexner and Brittney Ross for helpful comments on the manuscript. Research reported in this publication was supported by the NIDCD awards R01 DC013779 to MLF and F31DC016485 to JMR. The authors declare that the research was conducted in the absence of any commercial or financial relationships that could be construed as a potential conflict of interest.

## Methods & Materials

### Key Resources Table

| REAGENT or RESOURCE | SOURCE | IDENTIFIER |
| --- | --- | --- |
| Chemicals |  |  |
| Ketaset (ketamine) | Henry Schein Animal Health | Cat no. 010177 |
| AnaSed (xylazine) | Henry Schein Animal Health | Cat no. 033198 |
| Rimadyl (carprofen) | Henry Schein Animal Health | Cat no. 024751 |
| Ethyl valerate | Sigma-Aldrich | Cat no. 290866 |
| Ethyl butyrate | Sigma-Aldrich | Cat no. E15701 |
| Ethyl hexanoate | Sigma-Aldrich | Cat no. 148962 |
| Benzaldehyde | Sigma-Aldrich | Cat no. 418099 |
| 2-Heptanone | Sigma-Aldrich | Cat no. 537683 |
| Experimental Models:<br>Organisms/strain |  |  |
| Mouse: FVB/N-Tg(Thy1-cre)1Vln/J | The Jackson Laboratory | Stock no: 006143 |
| Mouse: B6;129S-Gt(ROSA)26Sortm38<br>(CAG-GCaMP3)Hze/J | The Jackson Laboratory | Stock no: 014538 |
| Software and<br>Algorithms |  |  |
| IDL v8.0 | Rubin-Tec | http://www.rubin-tec.co.il/ |
| Neuroplex v9.9.6 | RedShirtImaging, LLC | http://www.redshirtimaging.com/<br>support/soft_ov.html |
| ImageJ 1.49v | National Institutes of Health | https://imagej.nih.gov/ij/ |
| MATLAB R2015b | MathWorks | https://uk.mathworks.com/produ<br>cts/matlab.html |
| SPSS Statistics 22 | IBM | https://www.ibm.com/ms-en/marketplace/spss-statistics |
| R Studio v3.2.2 | RStudio, Inc | http://www.r-project.org |

#### Animals

All imaging and behavioral experiments used adult (8-14 weeks) male and female mice generated from crossing FVB/N-Tg(Thy1-cre)1Vln/J with B6;129S-Gt(ROSA)26Sortm38(CAG-GCaMP3)Hze/J, such that the resulting mice expressed the fluorescent calcium indicator GCaMP3 under the Thy1-promotor. Under this promoter, GCaMP3 is expressed in the olfactory bulb excitatory neurons, such as M/T cells (Chen et al., 2012). All experimental protocols were approved by the University of Tennessee Institutional Animal Care and Use Committee.

### General Methodology

#### Surgical Procedures

Mice were anesthetized with ketamine/xylazine (100/10 mg/kg, i.p.) and given analgesic injections (carprofen 10mg/kg s.c.) prior to surgery. Mice were secured in a custom stereotaxic apparatus (Narishige) and the bone overlying the dorsal surface of the olfactory bulb (OB) was thinned to create a cranial window for optical imaging. Additionally, some mice were implanted with bilateral cannula in the basolateral amygdala (BLA; Bregma: −1.5 AP, ± 3.3 ML, −5.0 DV). An anchor screw was inserted into the frontal or parietal bone and the entire skull was sealed with a thin layer of superglue (BSI). A custom made stainless steel headbar was attached to the posterior surface of each mouse’s skull for experimental head fixation during imaging. The entire skull was then covered with acrylic dental cement and mice were allowed two days to fully recover before experimentation.

#### Drug Infusions and placement verification

Mice with BLA cannula received bilateral intra-cannula muscimol (MUSC; 0.5μl of 0.5μg/μl delivered at a rate of 0.25μl/min) or an equal volume of vehicle (VEH; Ringers) infusions via a microsyringe pump. Injectors were left in place for an additional 2 minutes for diffusion. After completion of all experiments, mice with BLA cannula were perfused and brains were removed and sectioned to verify cannula placement within the BLA.

#### Optical Imaging

During imaging experiments, mice were head-fixed to a custom-built treadmill (Chettih et al., 2011; Heiney et al., 2014). The treadmill allowed mice to dictate their forward/reverse motions while remaining head-fixed. Imaging was performed using a Scientifica Slicescope equipped with a 4x objective (Olympus). The dorsal OB was illuminated with a LED light source centered at 480 nm. GCaMP3 signals were band-pass filtered with a Chroma emission filter (HQ535/50) and collected using a CCD camera at 25 Hz (NeuroCCD-SM256, Redshirt Imaging). During imaging experiments, mice experienced 5s trials consisting of 1s of no odor, followed by a 2s odor presentation, and 2s no odor (ITI= 1-2m). Odor presentations (ethyl valerate (E5), ethyl butyrate (E4), ethyl hexanoate (E6), benzaldehyde (BZ), 2-heptanone (2H)) were delivered to the nose via a flow-dilution olfactometer. Separate flow controllers for the clean air and the pure odorant vapor mixed the flow streams at the end of the odor delivery system to achieve an approximate concentration of 0.5% saturated vapor (s.v.) at a flow rate of 0.7 L/min.

### Behavior

#### Olfactory Fear Conditioning

All olfactory fear conditioning occurred in a standard shock chamber (Coulborn Instruments). Mice that underwent odor-shock conditioning (Paired) received 6 E5-foot shock pairings (10s E5 co-terminating with a 0.8mA, 0.5s foot shock). Any mice in the Shock only condition received 6 unpaired foot shocks of the same intensity, while mice in the Odor only condition received 6 unpaired E5 presentations of the same duration. Approximately 24 hours after conditioning, mice were placed in a novel, custom-made testing chamber. Mice were allowed a 5-10 minute acclimation period in the chamber before initiation of the testing protocol, in which we assessed behavioral freezing, a widely used measure of fear, for each of the 5 odors used during imaging experiments. This allowed testing of specific fear to the conditioned stimulus (E5) as well as generalized fear towards neutral odors (E4, E6, BZ, and 2H) never paired with shock. Testing consisted of one 20s presentation of each odor (ITI = 3 min), starting in the second minute. All testing odors were intensity matched and diluted in mineral oil to achieve an approximate headspace concentration of 200ppm. Odors were delivered to the testing chamber through dedicated lines.

#### Auditory Fear Conditioning

During fear conditioning, Tone-shock mice (n=3) were placed in the same shock chamber as olfactory fear conditioned mice but received 6 tone-shock pairings (10s 10kHz, 82dB, co-terminating with a 0.8mA, 0.5s foot shock). 24 hours later they were placed in the same novel context as olfactory conditioned mice but experienced 4-20s presentations of the paired tone to confirm tone-shock learning.

### Detailed methodology

*Experiment 1:* Mice underwent three consecutive days of chronic awake optical imaging (Pre1, Pre2, and Post) to assess glomerular odor representations for each odor. Following the Pre2 imaging session, mice were split into three groups for fear conditioning: Odor only, Shock only, and Paired, and then subjected to testing as detailed above (n = 5 each). Respiratory analysis (described below) were also conducted from these animals.

*Experiment 2:* Mice (n = 4) underwent two consecutive days of chronic awake imaging (Pre1 and Pre2) before fear conditioning to E5. Approximately 72 hours after paired conditioning, mice underwent testing and a final imaging session (Post3).

*Experiment 3:* Mice (n = 3) underwent the exact same experimental protocol as those in Experiment 1 except that all imaging sessions (Pre1, Pre2, and Post) were completed under anesthesia (100/10 mg/kg i.p. ketamine/xylazine).

*Experiment 4:* Mice (n = 3) underwent three consecutive days of chronic awake optical imaging, but only experienced E5 and BZ during imaging sessions. Following the Pre2 imaging session, mice were subjected to auditory fear conditioning and testing. The final, post-conditioning, imaging session was split into two halves (Post1 and Post2). During Post1, we assessed glomerular odor representations to E5 and BZ normally. During Post2, each odor trial was preceded by a 10s presentation of the same tone mice experienced during auditory fear conditioning to assess whether global fear states impact glomerular responses.

*Experiment 5:* During surgery, these mice received bilateral BLA cannula. Mice underwent the experimental procedures outlined in experiment 1 but received infusions of either VEH or MUSC (n = 5 each) 10 minutes before odor-shock conditioning to transiently inactivate the BLA during acquisition. During awake imaging, they experienced each of the five odors.

*Experiment 6:* During surgery, these mice received bilateral BLA cannula. Mice underwent three consecutive days of awake imaging, but only experienced E5, E4, and BZ during imaging. The final imaging session was split into two halves (Post1 and Post2), similar to Experiment 4. In this experiment, all mice were fear conditioned to E5 and tested for fear to each of the five odors before the final imaging session. Between Post1 and Post2, mice were left head-fixed on the treadmill and received infusions of either VEH or MUSC (n = 5 each) to transiently inactivate the BLA during expression. 10 minutes after the infusion, we resumed the Post2 imaging session half.

### Quantification and Statistical Analyses

Physiological data collected is based on number of responsive glomeruli and analysis of this data was achieved by collapsing data into a single value for each glomerulus representing its mean daily response. For visual clarity, graphical representation of this same data was reduced to a single value representing the mean daily glomerular response for each odor (i.e., five data points per mouse per day) and presented as mean ± SEM (standard error of the mean). Statistics were analyzed using R statistical analysis software and IBM SPSS 22.0. Equal variances were tested for all comparison data by Levene’s test or Mauchly’s test of Sphericity, and suitable corrections were made when necessary. Parametric statistical tests including t-tests, ANOVAs and Repeated Measures ANOVAs were performed, and post hoc analyses were conducted when main effects were found to be significant (Dunnett’s T for ANOVAs and Bonferroni with adjustment for multiple comparisons for RM ANOVAs).

Respiratory analyses were conducted using the raw fluorescent traces of each imaging trial. Raw fluorescent traces were picked from a random region of interest on the dorsal surface of the OB. Raw fluorescent traces were first smoothed by applying a rolling average of 3 frames across the entire trace. An algorithm was then used to detect each respiratory peak and to calculate the mean instantaneous frequency (MIF) between defined peaks before it was converted to respiration frequency (Hz). Odor-evoked MIF was restricted to the first four respirations after odor onset.

Spatial maps of stimulus-evoked glomerular activity were generated for each trial by first correcting for photobleaching as previously described (Fletcher et al., 2009). The odor-evoked change in fluorescence (ΔF) was then calculated by subtracting the average fluorescence of five frames centered around the peak of the respiration immediately preceding odor onset from five frames centered around the odor-evoked respiratory peak. Relative fluorescence change (ΔF/F) was then calculated by dividing the odor-evoked change in fluorescence (ΔF) by the average resting fluorescence gathered in the first 5 frames of the imaging trial. For spatial correlation analyses, the final normalized odor map for each odor was obtained by averaging all same-odor trials within each day to generate a daily mean odor map. Using ImageJ, daily mean odor maps were aligned across odors and days for individual animals. Next a pixel-by-pixel spatial correlation was performed to compare the daily mean odor map of each neutral odor to the map of E5 before (Pre2) and after (Post) training for the Paired, Odor only, and Shock only groups. For quantitative analysis of individual glomerular responses, trials were first separated by odor presentation. There was a minimum of 3 trials for each odor on each day. The individual glomerular responses for each trial were calculated from the ΔF/F measured at the center of each defined glomerulus (2x2 pixel average). Every individual responsive glomerulus was then normalized within each odor to its maximum observed ΔF/F response on Pre1 to that odor, such that the largest observed Pre1 response was equal to 1. A daily odor-evoked glomerular response was generated for each glomerulus by averaging all same-odor trials within a single day to allow for pooling across subjects and statistical comparison of pooled glomerular responses. Additionally, glomeruli were distinguished as E5 Responsive (i.e., glomeruli that respond to E5 alone or E5 and one or more of the other odors) or Non E5 Responsive (i.e., glomeruli that do not respond to E5) for further analysis.

For behavioral analysis, freezing was measured from the onset of each stimulus presentation for a total of 60 seconds. In Tone-Shock experiments the freezing of all four tone presentations was averaged for each mouse.

